# Gender- and gamete-specific patterns of X chromosome segregation in a three-gendered nematode

**DOI:** 10.1101/205211

**Authors:** Sophie Tandonnet, Maureen C. Farrell, Georgios D. Koutsovoulos, Mark L. Blaxter, Manish Parihar, Penny L. Sadler, Diane C. Shakes, Andre Pires-daSilva

## Abstract

Meiosis is at the core of sexual reproduction and alterations to its program can have dramatic effects. In this study, we investigate the segregation pattern of the X chromosome in *Auanema rhodensis*, a three-gendered nematode. This species has an atypical pattern of X chromosome segregation during male spermatogenesis that results in the exclusive production of haplo-X sperm. Here we use a combination of genetic and cytological approaches to show that while XX females undergo conventional meiosis to produce mostly haplo-X oocytes, hermaphrodites undergo atypical meiosis to produce nullo-X oocytes and mostly diplo-X sperm. Gender- and gamete-specific alterations of the normal meiotic program include non-pairing of the X homologs and precocious separation of X chromatids. Given these intra-species, intra-individual and intra-gametogenesis variations in meiotic program of *A. rhodensis*, we argue that it is an ideal model to study the plasticity of meiosis and how it can be modulated.

**Highlights:**

- Matings between *A. rhodensis* hermaphrodites and males generate only male cross-progeny.
- Hermaphrodites generate mostly nullo-X oocytes and diplo-X sperm.
- Following normal Mendelian genetics, XX females produce haplo-X oocytes.
- In cross-progeny, sons always inherit the X chromosome from the father.

## Introduction

In sexually reproducing organisms, meiosis is the process that enables diploid organisms to produce haploid gametes. This occurs in three key steps: (i) homologous chromosomes (“homologs”) pair and undergo crossovers that result in the molecular recombination of non-sister chromatids; (ii) homologs segregate to opposite poles; (iii) sister chromatids segregate to opposite poles. These meiotic processes not only ensure correct segregation of the genetic material, but also generates genetic variation through independent assortment and molecular recombination.

Errors in homolog pairing and/or recombination lead to subsequent meiotic defects, including non-disjunction between homologs and premature separation of sister chromatids during meiosis I (Zickler and Kleckner, 2015). The most common consequence of meiotic errors is aneuploidy, i.e. the abnormal number of chromosomes in the gamete. In humans, aneuploidy is the leading cause for miscarriages and developmental disorders such as Down’s (trisomy 21), Klinefelter (XYY) and Turner (XO) syndromes (Nagaoka et al., 2012).

The XX/XO sex determination system in the nematode *Caenorhabditis elegans* facilitates the study of meiosis because mutants are easy to recognize. Wild type hermaphrodites self-fertilize to give rise to mostly hermaphrodite progeny. In meiotic mutants, however, elevated aneuploidy of the X chromosome results in self-progeny with a high-frequency of males (Hodgkin, 1979, Meneely et al., 2002). The transparency of the animal, easy identification of the meiotic events along the gonad, simple karyotype (2*n* = 12), and genetic tools make *C. elegans* a powerful system to study meiosis (Rog and Dernburg, 2013).

The rhabditid nematode *Auanema rhodensis* (formerly called *Rhabditis* sp. SB347) (Kanzaki et al., 2017) offers the same advantages as *C. elegans* for studying meiosis: it has an XX/XO sex determination system, is small (~ 1 mm), free-living, has transparent gonad and can be easily maintained in the laboratory (Kanzaki et al., 2017, Shakes et al., 2011, Félix, 2004).

Furthermore, *C. elegans* provides a good reference for comparative studies for *A. rhodensis*, as they both are members of the Eurhabditis clade (Kiontke et al., 2007). However, unlike *C. elegans*, *A. rhodensis* simultaneously produce males, females and hermaphrodites (Félix, 2004, Kanzaki et al., 2017), a mating system called trioecy. Hermaphrodites and females are both XX and phenotypically almost indistinguishable, except for the fact that the former produces sperm in addition to oocytes (McCaig et al., 2017).

We previously reported that the *A. rhodensis* sex chromosome displays a non-canonical behavior during XO male gametogenesis (Shakes et al., 2011, Winter et al., 2017). The sister chromatids of the unpaired X chromosome prematurely separate during meiosis I. In addition, following anaphase II, an asymmetric partitioning of the cytoplasm occurs resulting in functional X-bearing sperm and nullo-X residual body (Shakes et al., 2011, Winter et al., 2017). This unique meiosis therefore systematically generates X-bearing sperm from XO males.

Here we report additional variations in the meiotic X chromosome segregation program of *A. rhodensis*. Notably, we find that the X chromosome segregation pattern differs between genders (females *versus* hermaphrodites) and gametogenesis type (oogenesis *versus* spermatogenesis) as well as within gametogenesis type. We discuss the genetic, population-genetic and evolutionary consequences of these modulations.

## Materials and Methods

### Strains

We used two isolates of *Auanema rhodensis*, originally derived from a deer tick (strain SB347, Rhode Island, USA) (Félix, 2004) and from a dead tiger beetle (strain TMG33, West Virginia, USA; found in May 2012, GPS 38.230011, −81.762252) (T. Grana, pers. commun.). Inbred strains were generated by picking single hermaphrodite animals from a self-fertilizing parent in every generation. The strain SB347, which underwent 50 generations of inbreeding, was subsequently renamed APS4. The strain TMG33, inbred for 11 generations, was renamed APS6. Strains were maintained at 20 °C according to standard conditions as for *C. elegans* (Stiernagle, 2006), either on MYOB agar (Church et al., 1995) for cytological studies or Nematode Growth Medium (Brenner, 1974) for molecular studies. Plates were seeded with the *Escherichia coli* uracil auxotroph mutant strain OP50-1. For molecular studies, microbial contamination was prevented by adding 200 μg/mL of nystatin and 200 μg/mL of streptomycin to the NGM.

### Genotyping of chromosomes

To genotype the X chromosome and autosomal linkage group 4 (LG4), we used 5 polymorphic markers (SNPs) for each chromosome (Table S1, Figure 4A). We generated these markers from a draft genome sequence for *A. rhodensis,* a genetic linkage map (in preparation) and strain-specific sequences (RAD-seq markers). The markers were selected for to the presence of a restriction enzyme site characteristic of one strain but not the other. Amplifications of the polymorphic regions were performed by single-worm PCRs followed by digestion of the products (Table S1). PCRs used the following conditions: 95 °C for 7 min, followed by 30-35 cycles of 15 s at 94 °C, 30 s at 55 °C, and 1 min at 72 °C. The digestion of the PCR products was performed at 35 °C for one to two hours. The genotype of each marker was visualized by gel electrophoresis of the digested products. The markers were confirmed to be X-linked by genotyping intra-species hybrid F1 males (XO). As expected from hemizygosity in XO animals, F1 males always showed a single genotype for markers on the X chromosome.

### Crosses between hermaphrodites and males

To distinguish hermaphrodite self-progeny from cross-progeny, we used morphologically-marked hermaphrodites (dumpy phenotype, strain APS19, caused by a recessive mutation). Ten crosses between a marked hermaphrodite and a wild type APS4 male were performed. The offspring were scored according to their phenotype (dumpy *versus* wild type) and gender at the adult stage. The female and hermaphroditic morphs were not distinguished and scored as “feminine”.

### Immunocytology

To obtain *A. rhodensis* adults of specific sexes, *A. rhodensis* hermaphrodites were isolated by selecting dauer larvae (Félix, 2004). Males and females were isolated from early broods of *A. rhodensis* hermaphrodites (Chaudhuri et al., 2015) and the gonads of females were secondarily verified by the absence of spermatogonia (McCaig et al., 2017).

To isolate meiotically dividing spermatocytes and meiotic one-cell embryos for analysis, hermaphrodites, males, mated females were dissected in Edgar’s buffer (Boyd et al., 1996) on ColorFrost Plus slides (Fisher Scientific) coated with poly-L-lysine (Sigma Aldrich Co.). Samples were freeze-cracked in liquid nitrogen and fixed in −20 °C methanol. Anti-tubulin labeling was done as previously described (Shakes et al., 2009) using 1:100 (0.025 mg/mL) FITC-conjugated anti-α-tubulin DM1A (Sigma-Aldrich). Slides were mounted with Fluoro-Gel II (Electron Microscopy Sciences) containing 6-diamidino-2-phenylindole (DAPI) and visualized under epi-illumination using an Olympus BX60 microscope.

## Results

### Genetic crosses suggest unorthodox patterns of meiotic X chromosome segregation that are both gender- and gamete-specific

Genetic crosses and cytological analyses show that *A. rhodensis* XO males produce exclusively haplo-X sperm (Shakes et al., 2011, Winter et al., 2017). Thus, crosses between males and females yield almost all feminine progeny (XX hermaphrodites or XX female) (Chaudhuri et al., 2015). Because the male sperm have a single X, this crossing result implies that most female oocytes carry a single X (Figure 1A). Without morphological genetic markers, it had been impossible to distinguish between self and outcross progeny from hermaphrodite oocytes fertilized by male sperm. Using our new, morphologically-marked strain containing a recessive dumpy mutation, we performed crosses between dumpy hermaphrodites and wild type males. In such crosses, cross-progeny were easily distinguished by their non-dumpy phenotype. Contrary to our expectations, all cross-progeny were male (306 normal males scored from 10 hermaphrodite/male crosses).

Since male sperm have a single X, this result implies that XX hermaphrodites produce oocytes without an X (nullo-X oocytes) (Figure 1B). In turn, as self-fertilizing hermaphrodites produce 90-95% XX feminine progeny (Félix, 2004, Chaudhuri et al., 2015, Farrell, 2015), their nullo-X oocytes must be fertilized by hermaphrodite sperm that are predominantly diplo-X (Figure 1C).

**Figure 1.**
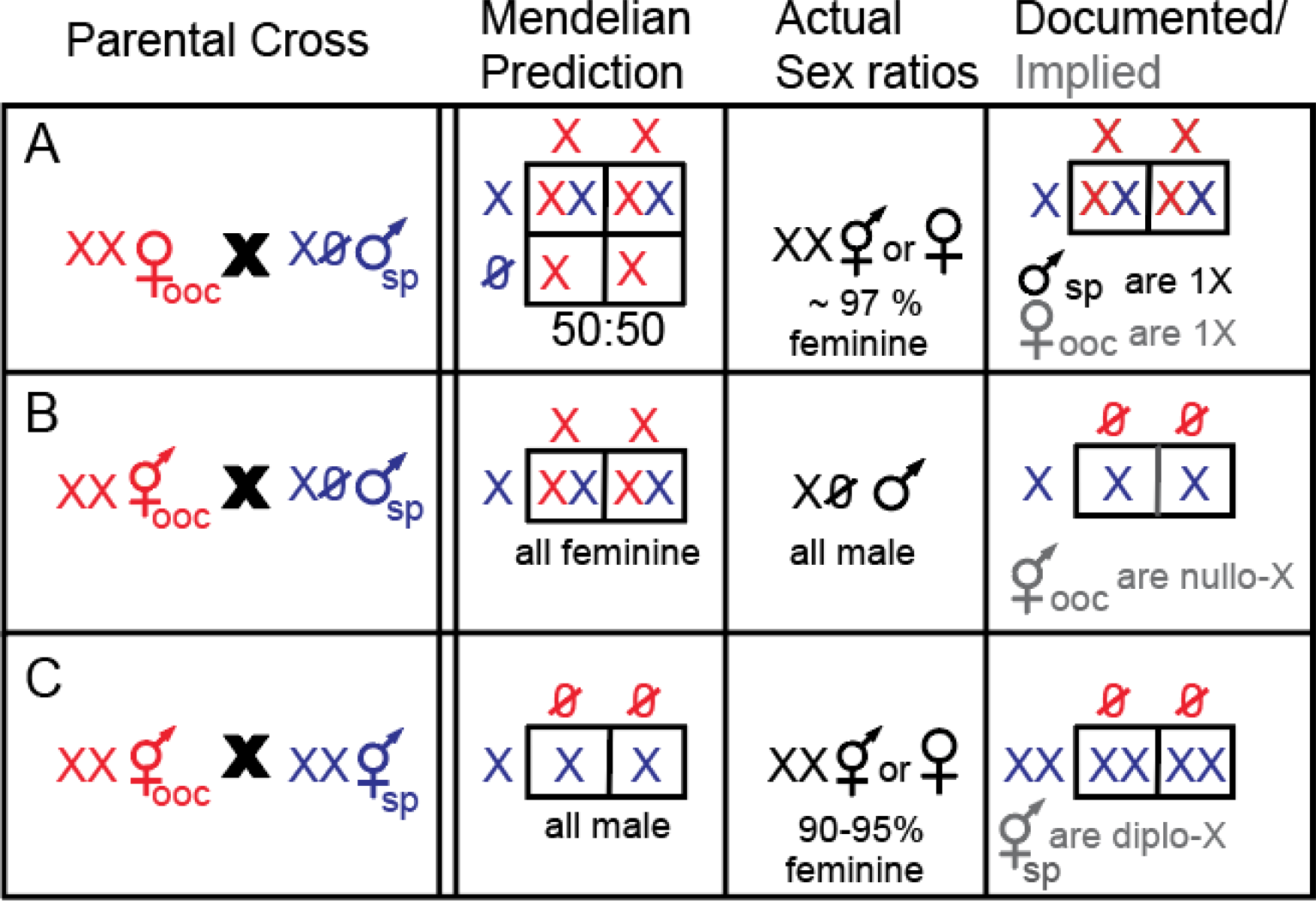
Schematic of genetic crosses and implied patterns of X chromosome segregation in *A. rhodensis*. **(A)** Crosses between XX females and XO males generate mostly XX progeny, since males mainly produce haplo-X sperm. This result implies that oocytes are haplo-X. **(B)** Crosses between XX hermaphrodites and XO males result only in male progeny, implying the production of nullo-X oocytes by females. **(C)** Self-fertilization of an XX hermaphrodite results mostly in XX progeny, implying that sperm are diplo-X. In red are the gametes produced during oogenesis (ooc) and in blue gametes produced during spermatogenesis (sp).

### Cytological analysis of meiotic X chromosome segregation

#### Oogenesis Patterns

Our crossing results predicted specific cytological consequences. We hypothesized that during oogenesis in *A. rhodensis* hermaphrodites, unorthodox segregation patterns of the X chromosome would result not only in anaphase figures with unequal amounts of chromatin, but also in non-standard numbers of DAPI-stained bodies aligned at the metaphase plate due to potential alterations in X chromosome pairing. We examined meiotically dividing oocytes labelled with a combination of DAPI-staining and anti-tubulin antibodies and compared the patterns in *A. rhodensis* females and hermaphrodites to the well-established patterns in *C. elegans* (Albertson and Thomson, 1993, Golden et al., 2000, Dumont et al., 2010, Cortes et al., 2015).

During *C. elegans* oogenesis, chromosome condensation occurs over an extended period during late meiotic prophase (Albertson and Thomson, 1993). Thus, it is relatively easy to observe metaphase I figures with six bivalents (five autosomal and one X). In contrast, chromosome condensation in *A. rhodensis* occurs rapidly between the end of meiotic prophase and metaphase I (data not shown), and thus metaphase I figures with well-resolved chromosomes were relatively rare. When we did observe them, the metaphase I figures in *A. rhodensis* females had seven DAPI-stained structures, consistent with genomic analyses that suggest *A. rhodensis* has six autosomes and an X (unpublished). In the oocytes of *A. rhodensis* females, chromosome segregation patterns during both anaphase I and anaphase II were equal, although we did find examples of lagging chromosomes during early anaphase I (Table 1). In contrast, analyses of hermaphrodite oocytes in *A. rhodensis* revealed two key differences. First, the metaphase I figures had either seven or eight DAPI-stained structures. Observing eight structures is consistent with the presence of X chromosomes that had failed to pair or recombine. Second, anaphase I figures typically exhibited either lagging chromosomes or unequal chromosome segregation (Figure 2; Table 1). Also consistent with the unequal pattern of chromosome segregation, the first polar bodies were proportionally larger, while anaphase II figures were consistently equal. Taken together, the frequent observation of an additional DAPI staining body in metaphase I plates of *A. rhodensis* hermaphrodite oocytes and the unequal divisions observed during anaphase I, suggest a model in which the X chromosomes of hermaphrodite oocytes fail to pair and/or recombine during meiotic prophase and then are partitioned to the first polar body during anaphase I.

**Figure 2.**
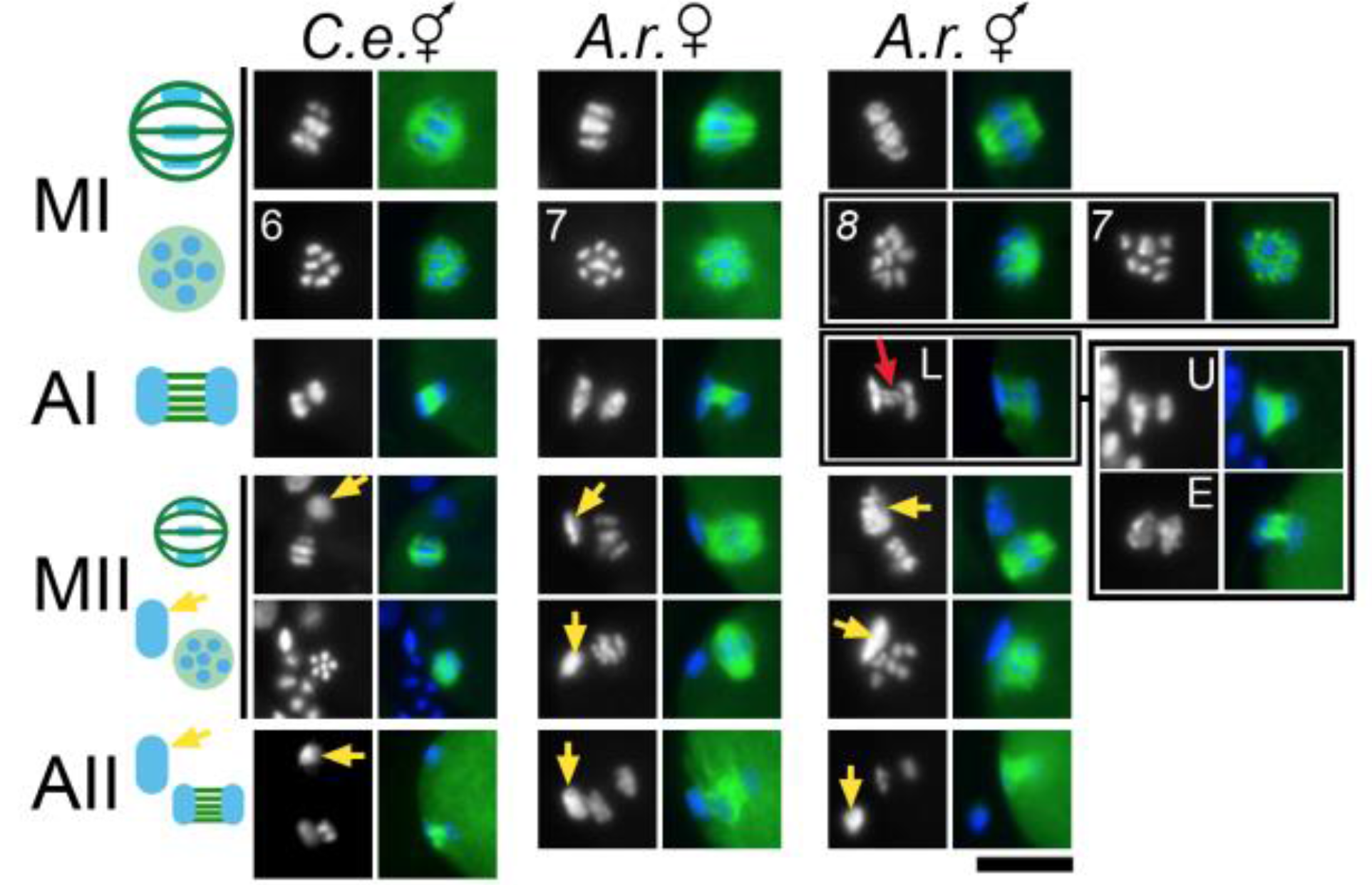
Patterns of chromosome segregation during oocyte meiosis. Chromosome segregation patterns were imaged in fixed, meiotic one-cell embryos. Chromosomes were stained with DAPI (blue) and microtubules were labelled with anti-tubulin antibody (green). Schematics of the meiotic divisions are shown in the left column. Metaphase spindles are shown in two orientations; either from the side (upper) or viewed down the pole to show the metaphase plate. For *A. rhodensis* (*A. r.*) hermaphrodites, metaphase I plates with 8 and 7 DAPI-staining bodies are shown as well as anaphase I figures with lagging chromosomes (L), unequal chromosome segregation (U), and a rare example of an equal chromosome segregation (E). The red arrow shows lagging chromosomes during anaphase I in *A. rhodensis* hermaphrodites. The yellow arrows show polar bodies. Abbreviations: metaphase I (MI); anaphase I (AI), metaphase II (MII), and anaphase II (AII). Scale bar = 10 microns.

**Table 1.** Scoring of anaphase figures during oogenesis.

| Species and gender | Stage of oocyte meiosis | Lagging Chromosome | Equal Segregation | Unequal Segregation |
| --- | --- | --- | --- | --- |
| *C. elegans* hermaphrodite | Anaphase I (n=15) | 0% | 100% | 0% |
|  | Anaphase II (n=6) | 0% | 100% | 0% |
| *A. rhodensis* female | Anaphase I (n=28) | 10% | 90% | 0% |
|  | Anaphase II (n=3) | 0% | 100% | 0% |
| *A. rhodensis* hermaphrodite | Anaphase I (n=46) | 24% | 6% | 70% |
|  | Anaphase II (n=29) | 0% | 100% | 0% |

#### Spermatogenesis patterns

Previously we showed that sperm production in *A. rhodensis* hermaphrodites differs from that in *C. elegans,* since *A. rhodensis* hermaphrodites produce sperm from distinct clusters of spermatogonial cells - both simultaneously and continuously along with oocytes (McCaig et al., 2017). In addition, *A. rhodensis* hermaphrodites, like *A. rhodensis* males, produce only two rather than four functional sperm during meiosis. We have previously assumed that hermaphrodite sperm, like those in *A. rhodensis* males, contained a single X (Winter et al., 2017, Shakes et al., 2011). However, if *A. rhodensis* hermaphrodites are routinely producing nullo-X oocytes, the production of predominantly XX progeny by self-fertilizing hermaphrodites predicts that XX hermaphrodites are not making haplo-X sperm, but rather diplo-X sperm. To test this prediction, we examined meiotically dividing spermatocytes in *A. rhodensis* hermaphrodites and compared them with patterns that previously described in males (Winter et al., 2017, Shakes et al., 2011).

In *A. rhodensis* XO male spermatocytes, the X chromatids precociously separate in meiosis I, resulting in X chromosomes segregating symmetrically (no lagging X is observed) (Shakes et al., 2011). During anaphase II, the lagging X chromatid invariably ends up in the functional male sperm, whereas the other chromosomal complement is discarded in a ‘residual body’ (Figure 3; (Winter et al., 2017, Shakes et al., 2011)). In XX hermaphrodite spermatocytes, the segregation patterns were similar. Analysis of 520 hermaphrodite gonads yielded 16 clusters with anaphase II stage spermatocytes, and the cells within all 16 clusters exhibited a lagging, potentially unresolved, DAPI-staining chromatin mass that was roughly twice the size of those in anaphase II male spermatocytes (Figure 3). In the same set of specimens we identified 17 clusters with post-meiotic, partitioning stage sperm, and in each case the functional sperm appeared to have more DNA than the tubulin-containing residual body (Figure 3). These observations, taken together, provide cytological evidence that the hermaphroditic sperm likely contain two X chromosomes.

**Figure 3.**
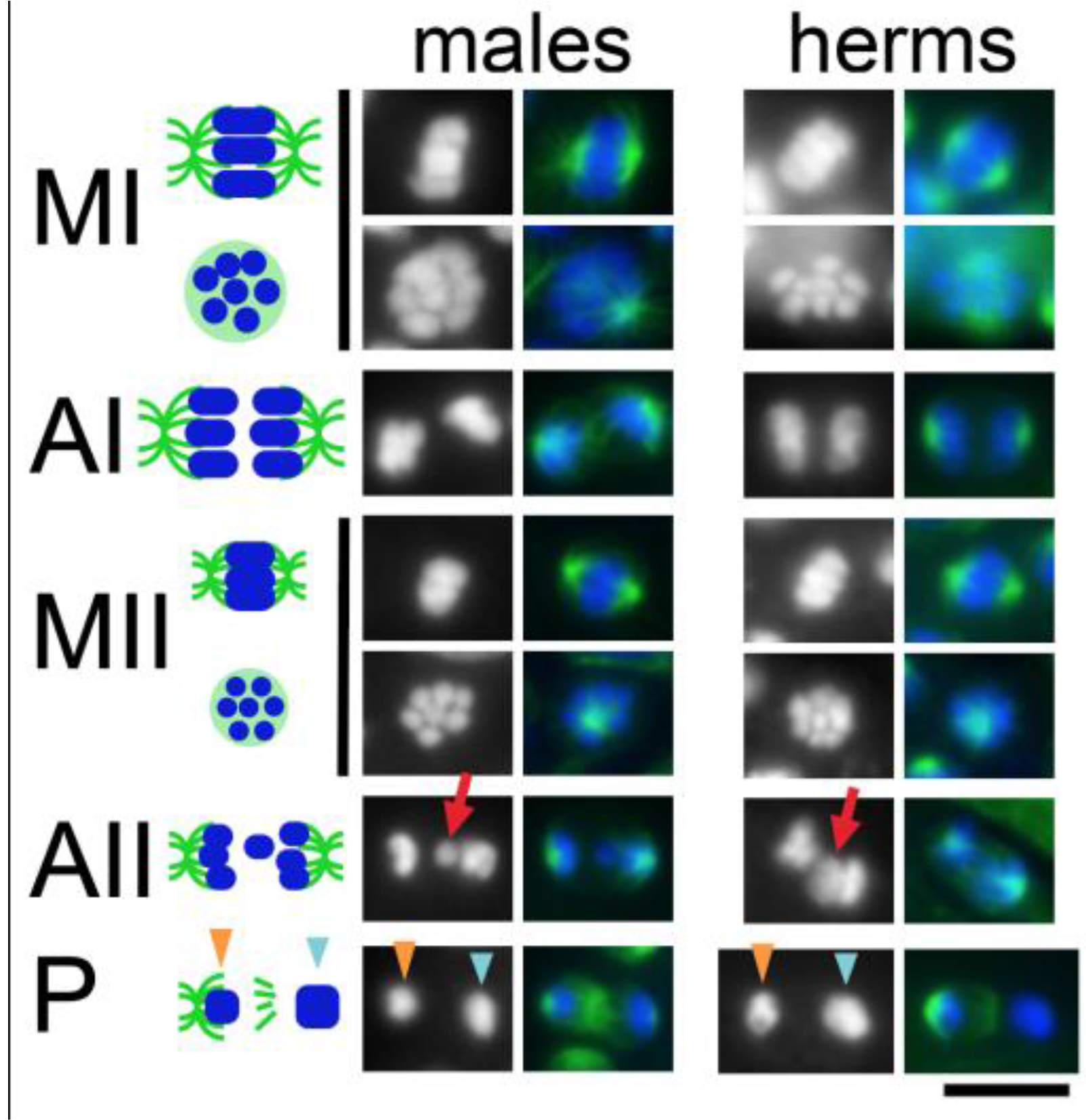
Patterns of chromosome segregation during *A. rhodensis* spermatocyte meiosis. Chromosome segregation patterns were imaged in isolated and fixed male and hermaphrodite gonads. Chromosomes were stained with DAPI (blue) and microtubules were labelled with antitubulin antibody (green). A schematic of the meiotic divisions are shown in the left column. Metaphase spindles are shown in two orientations to either show the spindle, or viewed down the pole to show the metaphase plate. The red arrows indicate lagging chromosomes during anaphase II. The orange arrowheads indicate the chromatin mass of the future residual body during anaphase II and the partitioning (P) phase. The blue arrowheads indicate the larger chromatin mass of the future sperm. Meiotic stage abbreviation as in Figure 2. Scale bar = 5 microns.

### Independent analysis of X chromosome segregation patterns using strain specific SNP markers

To track segregation patterns of X chromosomes from maternal and paternal parents, we identified single nucleotide polymorphisms (SNPs) between two independently isolated strains of *A. rhodensis* (APS4 and APS6). Using 5 polymorphic markers distributed along the length of the X chromosome (Figure 4A), we followed the pattern of inheritance of the X chromosome in F2 individuals produced by hybrid (X_APS4_X_APS6_) female or hermaphrodite mothers derived from original crosses between the inbred strains APS4 (carrying X_APS4_) and APS6 (carrying X_APS6_)(Figure 4B).

#### X chromosomes in females

Intra-specific hybrid (X_APS4_X_APS6_) F1 females crossed with males of one of the parental strains (e.g. X_APS6_) produced F2 feminine progeny with the expected 1:1 ratio of homozygous (X_APS6_X_APS6_) to heterozygous (X_APS6_X_APS4_) markers in the X chromosome (chi-squared = 3.37, df = 1, p-value = 0.07, Table 2, Table S2). Notably, we also found 12 examples of crossover, where the X genotyping showed some markers as heterozygous and others as homozygous in a same individual (Figure 4C, Table 2, Figure S1 and Table S2). These data point towards a conventional meiotic pairing and segregation of the X chromosome in *A. rhodensis* females.

**Table 2.**
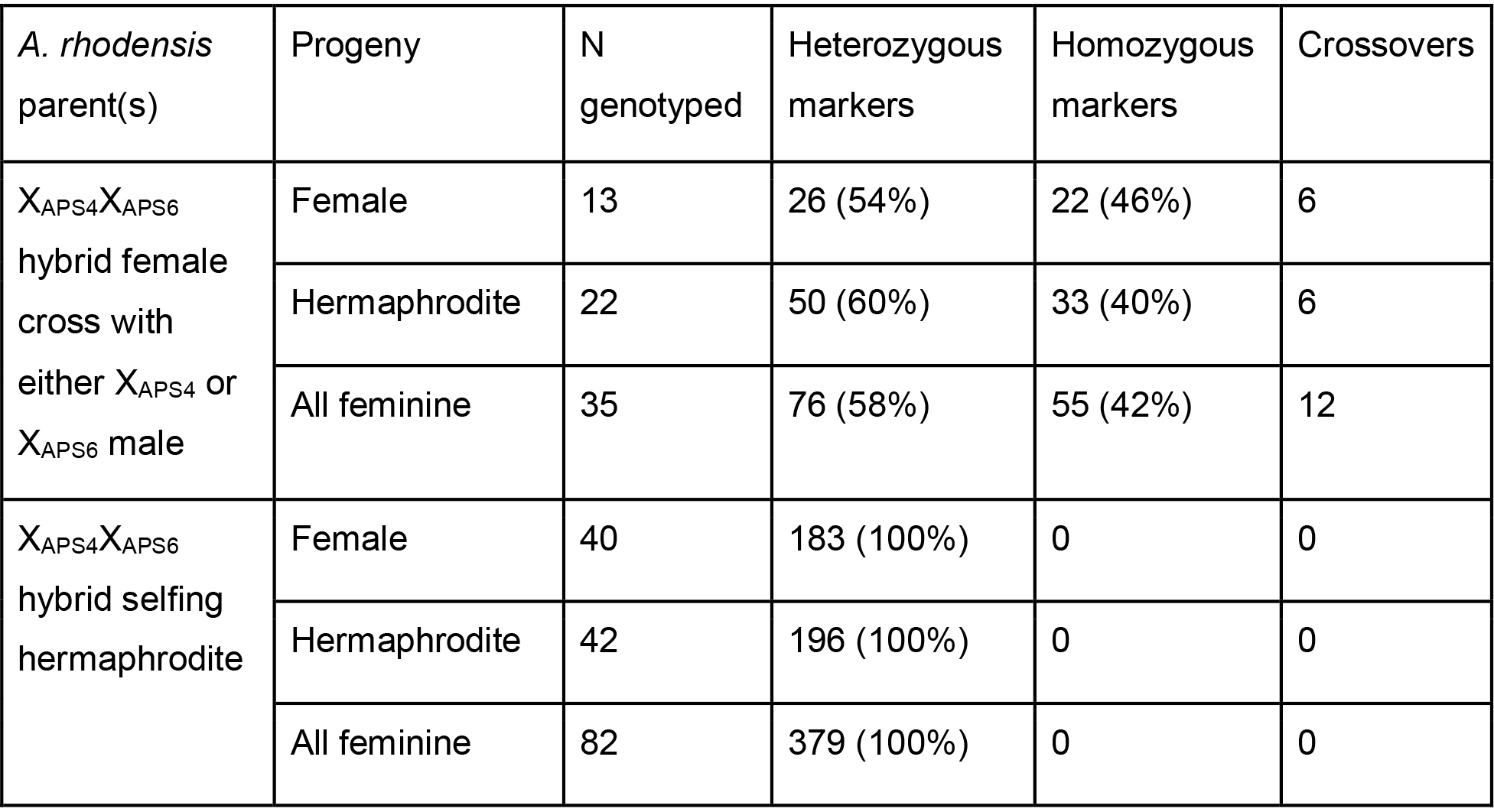
Genotype counts of F2 feminine progeny (female and hermaphrodite) from hybrid F1 crossed female or hybrid F1 selfing hermaphrodite mothers. It was not always possible to visualize crossovers as the genotyping procedure failed for some markers in several individuals and that only 5 markers were used.

| *A. rhodensis*<br>parent(s) | Progeny | N<br>genotyped | Heterozygous<br>markers | Homozygous<br>markers | Crossovers |
| --- | --- | --- | --- | --- | --- |
| $X_{APS4}X_{APS6}$<br>hybrid female<br>cross with<br>either $X_{APS4}$ or<br>$X_{APS6}$ male | Female | 13 | 26 (54%) | 22 (46%) | 6 |
|  | Hermaphrodite | 22 | 50 (60%) | 33 (40%) | 6 |
|  | All feminine | 35 | 76 (58%) | 55 (42%) | 12 |
| $X_{APS4}X_{APS6}$<br>hybrid selfing<br>hermaphrodite | Female | 40 | 183 (100%) | 0 | 0 |
|  | Hermaphrodite | 42 | 196 (100%) | 0 | 0 |
|  | All feminine | 82 | 379 (100%) | 0 | 0 |

**Figure 4.**
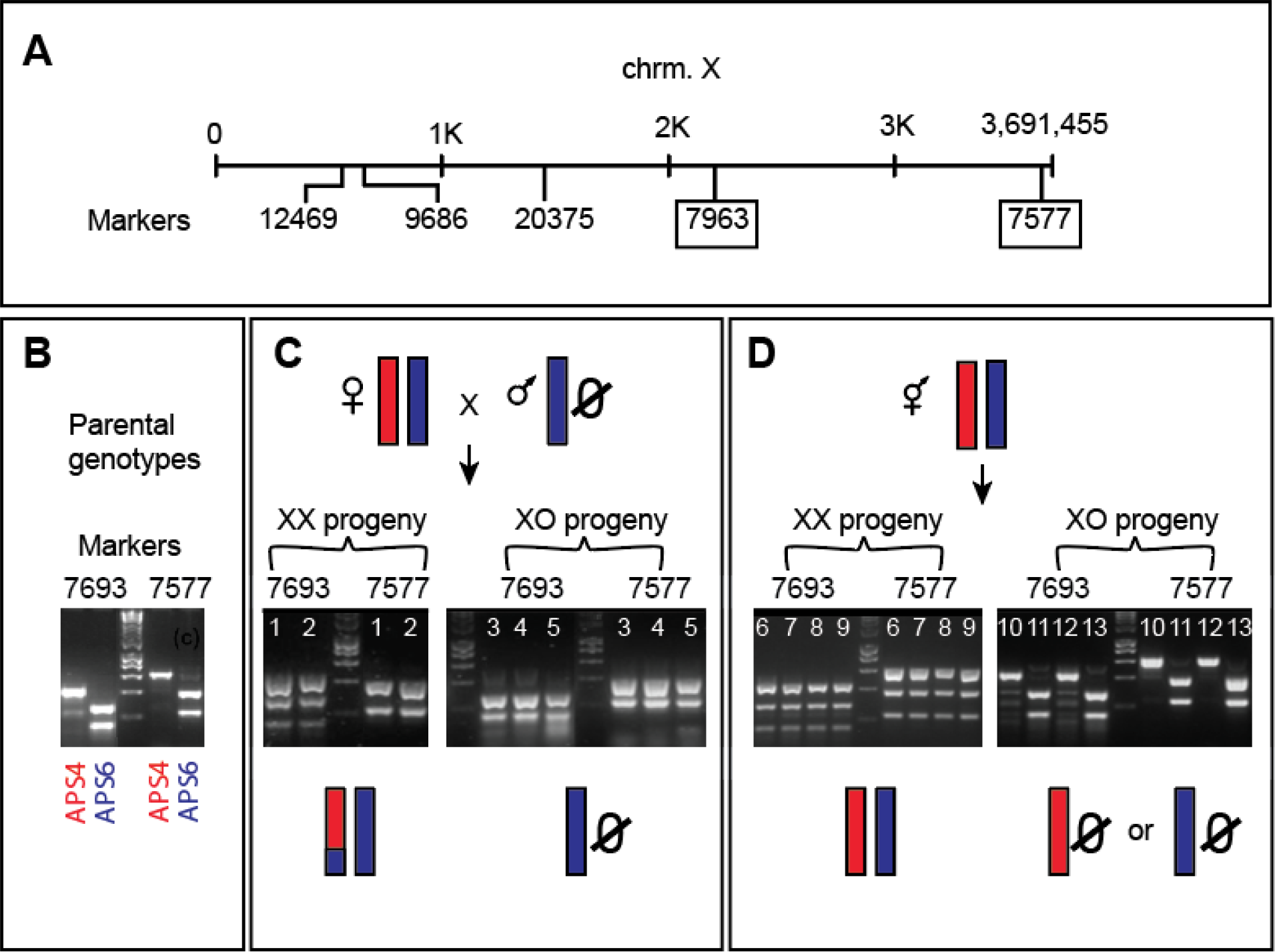
Genotyping markers and representative example of the X genotyping results. **(A)** Schematic view of the markers used to genotype the X chromosome. **(B-D)** Illustration of the inheritance pattern and representative genotyping profiles of the X chromosome for a pair of markers (7963 and 7577). **(B)** Genotyping profile of parental strains. **(C)** A hybrid female crossing with an APS6 male results in XX progeny with both homozygous and heterozygous X markers. Crossovers could be detected when the X of one individual was part heterozygous, part homozygous, as represented here by individuals 1 and 2. Male offspring resulting from the cross always inherited the X from their father. **(D)** X genotyping of individuals produced by hybrid selfing hermaphrodites reveals that the X chromosome remains heterozygous in XX individuals and hemizygous for each parental strain in males. Numbers in each gel lane represent individual animals.

#### X chromosomes in hermaphrodites

Analysis of F2 feminine progeny produced by selfing F1 (X_APS4_X_APS6_) hermaphrodites revealed a very different pattern. These F2 feminine progeny were invariably heterozygous for the X chromosome (Figure 4D, Table 2, Figure S2, Table S2). For this analysis, 82 F2 feminine progeny produced by X_APS4_X_APS6_ F1 hybrid hermaphrodites were genotyped across 5 markers positioned along the X chromosome (Figure 4A). Although Mendelian segregation patterns would predict a 1:2:1 ratio of X_APS4_X_APS4_: X_APS4_X_APS6_: X_APS6_X_APS6_ progeny amongst the feminine F2s, all individuals were heterozygous (X_APS4_X_APS6_) across the 379 markers successfully genotyped (Figure 4D, Table 2, Table S2), which implies that no recombination between the X chromosomes took place. Given that we have shown that the feminine progeny of selfing hermaphrodites inherit both of their X chromosomes from the diplo-X hermaphrodite sperm, this SNP analysis shows that the two X chromosomes in the diplo-X sperm are homologs, not sisters. This X chromosome behavior is consistent with a model in which both X chromosomes of a hermaphrodite spermatocyte separate into sister chromatids in meiosis I and then both X chromatids segregate to the functional sperm in meiosis II (Figure 5D).

Importantly, this behavior was specific to the X chromosome, as genotyping of the autosome LG4, also across 5 markers (Table S1,Figure S3), yielded a mix of homozygous and heterozygous markers in keeping with Mendelian expectations (24 and 17, respectively). In addition, autosomal crossovers and double crossovers could be observed, as the genotype was not uniform across all markers for a same individual (Table S2, second page).

**Figure 5.**
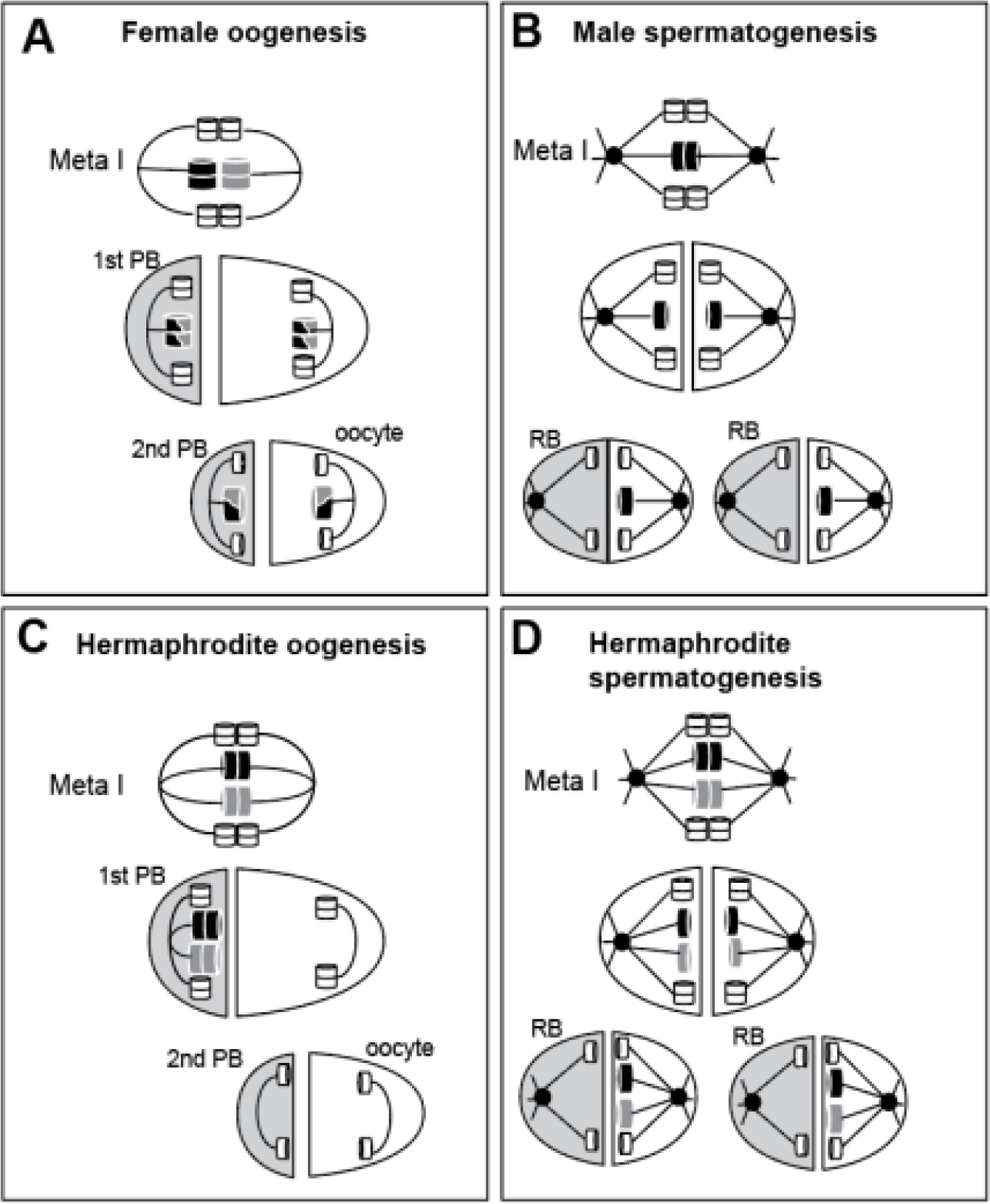
Simplified model of the X chromosome segregation mechanism in *A. rhodensis*. **(A)** In females, autosomes (white cylinders) and X chromosomes (darker and larger cylinders) dynamics follow the canonical segregation, with pairing and crossover. Shaded cells are polar bodies (PB). Lines represent the microtubules. **(B)** In XO males, the homologous autosomes segregate to different daughter cells in meiosis I, and sister chromatids separate in meiosis II. For the unpaired X chromosome, however, sister chromatids separate in meiosis I. X-bearing cells become spermatids, whereas nullo-X cells are discarded into residuals bodies (RB, shaded in grey). Black circles represent centrioles. **(C)** During meiosis I of hermaphrodite oogenesis, non-disjunction of the X chromosomes occurs, leading to diplo-X polar bodies in meiosis I and nullo-X functional oocytes after meiosis II. **(D)** In hermaphrodite spermatogenesis, the homologous X chromosomes fail to pair and the sister chromatids separate during meiosis I, similarly to the male spermatogenesis **(A)**. Only X-bearing cells become functional spermatids.

### Father-to-son X chromosome inheritance

We next asked how males are generated in populations of *A. rhodensis*. To do so, we contrasted the canonical meiosis of females with the atypical one of hermaphrodites (asymmetric distribution of the X chromosomes). *A. rhodensis* populations are comprised of 5-10% males (XO), which can be generated either from selfing hermaphrodites or from female/male crosses (Félix, 2004, Chaudhuri et al., 2015).

We genotyped the X chromosomes of males produced by female/male crosses or by selfing hermaphrodites. Sons resulting from female/male crosses always inherited the X markers of their father (Table 3, Table S2). The production of male progeny from crosses between males and females implies that unusual meiotic divisions must sometimes occur during female oogenesis, generating nullo-X oocytes. The rare production of nullo-X female oocytes is likely to be mechanistically similar to the routine production of nullo-X hermaphrodite oocytes.

To explain the occurrence of male offspring from selfing hermaphrodites, we postulate that hermaphrodite spermatocytes sometimes divide to generate haplo-X rather than diplo-X sperm. Intriguingly, selfing hermaphrodites regularly produce more males early in their broods (Chaudhuri et al., 2015), suggesting that the choice of the division pattern is developmentally regulated. This mode of division does not seem to require crossovers, as the X chromosome of males from selfing X_APS4_X_APS6_ hermaphrodites was either entirely X_APS4_ or X_APS6_ for all the markers genotyped for a same individual (100 genotypes from 21 males, Figure 4D, Table S2).

**Table 3.** X chromosome genotyping of F1 males resulting from crosses between the APS4 and APS6 parental strains.

|  | **APS4 ♀ x APS6 ♂ cross** |  | **APS6 ♀ x APS4 ♂ cross** |  |
| --- | --- | --- | --- | --- |
| **F1 male genotype** | **APS4** | **APS6** | **APS4** | **APS6** |
| Marker 9686 | 0 | 21 | 18 | 0 |
| Marker 12469 | 0 | 23 | 18 | 0 |
| Marker 20375 | 0 | 20 | 20 | 0 |
| Marker 7963 | 0 | 20 | 15 | 0 |
| Marker 7577 | 0 | 18 | 19 | 0 |

## Discussion

We have shown that meiosis of the sex chromosome can be modulated within a species in an identical genetic context. *A. rhodensis* is unusual in having three genders: male, female and hermaphrodite. The meiosis program governing X chromosome inheritance in this nematode varies with the gender of the parent, the gametogenesis type and even within the same gametogenesis. Female oogenesis displayed a classical meiosis with recombination and Mendelian segregation of the X chromosomes, generating the expected haplo-X gametes (Figure 5A). In both hermaphrodite oogenesis and spermatogenesis, the X chromosomes fail to pair. In hermaphrodite oogenesis, the X chromosomes segregated to the same pole (first polar body), yielding nullo-X oocytes (Figure 5C). During hermaphrodite spermatogenesis, a precocious sister chromatid separation of the X chromosomes during meiosis I followed by a partitioning of the X chromatids to the functional sperm during meiosis II, led to the production of diplo-X sperm similarly of the dynamics observed during male spermatogenesis. One consequence of this division is that the two X chromatids contained in the hermaphrodite sperm are homologs and not sisters.

X chromosome fates in hermaphrodite oogenesis and spermatogenesis balance each other, as diplo-X sperm fertilize nullo-X oocytes to create XX zygotes. A direct outcome of this atypical system is that it maintains the heterozygosity of X chromosomes transmitted during hermaphrodite reproduction, as crossovers and thus recombination between the X homologs never occurs. In this manner, X chromosome segregation differs between gametogenesis in hermaphrodites compared to females, and between spermatogenesis and oogenesis within the same hermaphrodite. As genetically identical X chromosomes can be segregated differentially within each individual, control of the atypical meiosis process observed in hermaphrodite gametogenesis cannot lie in the X chromosome sequence per se.

Even though the molecular mechanisms mediating the recognition of homologs are still not fully understood, chromosomal regions named pairing centers (PCs) play a major role in the pairing, synapsis, crossover and disjunction of homologs in *C. elegans* (Rog and Dernburg, 2013, MacQueen et al., 2005). The recruitment of various proteins including ZIMs/HIM-8, PLK-2, SUN-1, and ZYG-12 at the site of the PCs is required to promote and stabilize homolog pairing and to link the PCs to microtubules (reviewed in (Rog and Dernburg, 2013)). The zinc-finger protein HIM-8 specifically targets the X chromosome PCs of *C. elegans* and is required for proper X chromosome segregation (Phillips et al., 2005). In *him-8* mutants, the X chromosome homologs fail to pair and synapse, which results in a high rate of X non-disjunction. In *A. rhodensis*, where there is a lack of pairing between the X homologs during hermaphrodite oogenesis, it may be that *trans-acting* factors specific to X chromosome meiosis, such as HIM-8, are differentially regulated between female and hermaphrodite oogenesis. As observed in *C. elegans* (Cortes et al., 2015), the resulting X univalents would be preferentially placed in the first polar body and thus eliminated.

In *C. elegans*, the cis-acting *me8* mutation directly alters the PCs and results in a lack of crossovers and disjunction of the X chromosome (Villeneuve, 1994). In *A. rhodensis*, because the modulations of X chromosome segregation are independent of the sequence of the chromosome, this genetic mechanism cannot be the cause of the variant segregation behaviors. However, the X chromosome (particularly the PC region) could be subject to differential chromatin modification that affects homologous pairing or recombination. Heteromorphic sex chromosomes have been shown to undergo more condensation than autosomes. This is thought to prevent harmful non-homologous recombination between heterogametic chromosomes (McKee and Handel, 1993). In accord with these findings, the X chromosome of *C. elegans* appears highly condensed and repressed during male meiosis (Kelly et al., 2002). Despite the fact that the X chromosomes of hermaphrodites are, surprisingly, also found compacted during meiosis, the pairing and recombination between X homologs follows a different pattern than that of the autosomes (Kelly et al., 2002), suggesting the presence of an X-specific meiotic machinery.

The premature separation of sister chromatids (PSSC) of the X chromosomes observed during the spermatogenesis of *A. rhodensis* hermaphrodites is reminiscent of the atypical X chromatid separation occurring during meiosis I of male spermatogenesis (Shakes et al., 2011, Winter et al., 2017). The X PSSC in both male and hermaphrodite spermatogenesis could be governed by the same mechanism. The cohesin complexes, and particularly the kleisin subunits REC-8 and COH-3/4, necessary to hold the sister chromatids together, could be involved as they have been shown to play an important role of the correct segregation of homologs (reduction of ploidy) and chromatids in *C. elegans* (Severson and Meyer, 2014, Pasierbek et al., 2001). Indeed, REC-8 and COH-3/4 cohesins are crucial for the efficient formation of crossovers and correct assembly of the synaptomemal complex between homologs (Severson et al., 2009, Severson and Meyer, 2014). REC-8 in particular helps the co-orientation of the sister chromatids towards the same pole (Severson et al., 2009, Severson and Meyer, 2014). Mutations of the *rec-8* gene induce premature separation of the sister chromatids and lack of connection between homologs (Pasierbek et al., 2001). In *A. rhodensis* it is possible that cohesin complexes required to tether the sister chromatids during meiosis I follow a different dynamic, allowing the precocious separation of the X chromatids during spermatogenesis diakinesis.

In male and hermaphrodite spermatogenesis (Shakes et al., 2011, Winter et al., 2017), this study), the X chromosomes segregate to the pole fated to become functional sperm. A remarkable consequence of this system is that the X chromosome is inherited through the sperm-producing germline and from father to son in the event of a cross. As far as we know, this is the only example of a complete X chromosome transmission through the male lineage. One evolutionary consequence of this observation is that any beneficial mutations on the X will spread very quickly through the population as male carriers will transmit it to all their offspring including their sons, which will, in turn, systematically pass it on. Additionally, as there is no crossover between X chromosomes during hermaphrodite production of XX offspring, this means that the *A. rhodensis* X chromosome has a very different recombinational and evolutionary trajectory than does the *C. elegans* X. If X-linked genes control traits subject to selection, the maintenance of diversity in X chromosomes in feminine nematode offspring of hermaphrodites could impact the colonizing ability of single hermaphrodite nematodes.

The unusual inheritance of the X through the paternal lineage implies that the feminine germlines must produce nullo-X oocytes. We have shown that during hermaphrodite oogenesis, nullo-X oocytes are prevalent. However, female oogenesis usually follows a conventional Mendelian segregation of the X chromosome resulting in haplo-X oocytes. Thus an atypical segregation of the X must occasionally occur during female oogenesis. Selfing hermaphrodites also produce a small percentage of males (~5-10%) (Chaudhuri et al., 2015) and therefore must occasionally, produce haplo-X gametes. Because sperm are produced in spermatogonial clusters within the hermaphrodite germline (McCaig et al., 2017), it may be that different clusters produce sperm with different X chromosome complements. These observations indicate that the meiosis program is actively modulated within the same gametogenesis, generating a flexible system where the proportion of male offspring can be adjusted through regulation of the X chromosome segregation in both female and hermaphrodite mothers. The factors controlling this regulation, and thus the male:feminine sex ratio, could be environmental and may reflect adaptation to the colonization ecology of *A. rhodensis*.

The recent findings and data collected on *A. rhodensis* open the door to investigate the peculiarities and implications of its sex determination system, understand mechanistically the processes that control X chromosome segregation, and explore the evolutionary and population genetic consequences of the curious pattern of X chromosome inheritance. *A. rhodensis* is mutable, and screening for genetic loci that specifically affect female, hermaphrodite or male X chromosome segregation (i.e. the proportion of male offspring generated) is feasible given the genetic and genomic resources we have generated. Particularly, *A. rhodensis* is an ideal model to study the regulation of the meiosis process and how it can be altered within a same genetic context. We note that developmental context (hermaphrodite *versus* female) plays an important role in the modulation of meiotic processes affecting the X. For instance, XX animals that develop through a dauer larva stage always become hermaphrodites (Chaudhuri et al., 2011), whereas larvae that bypass this stage become females. What triggers this differential development and how it links with the meiotic process are still open questions.

## Acknowledgments

G.D.K. was supported by a BBSRC PhD studentship and S.T. by a PhD training grant from CAPES/CNPq (201116/2014-6). A.P.S. was supported by a grant from National Science Foundation (IOS 1122095), BBSRC (BB/L019884/1) and University of Warwick start-up funds. D.C.S. was supported by a grant from the National Science Foundation (IOS 1122101). We thank Dr Sally Adams, Dr Pedro Robles Naharro and Dr James Stratford for many insightful conversations.

## Supplementary material

**Table S1.**
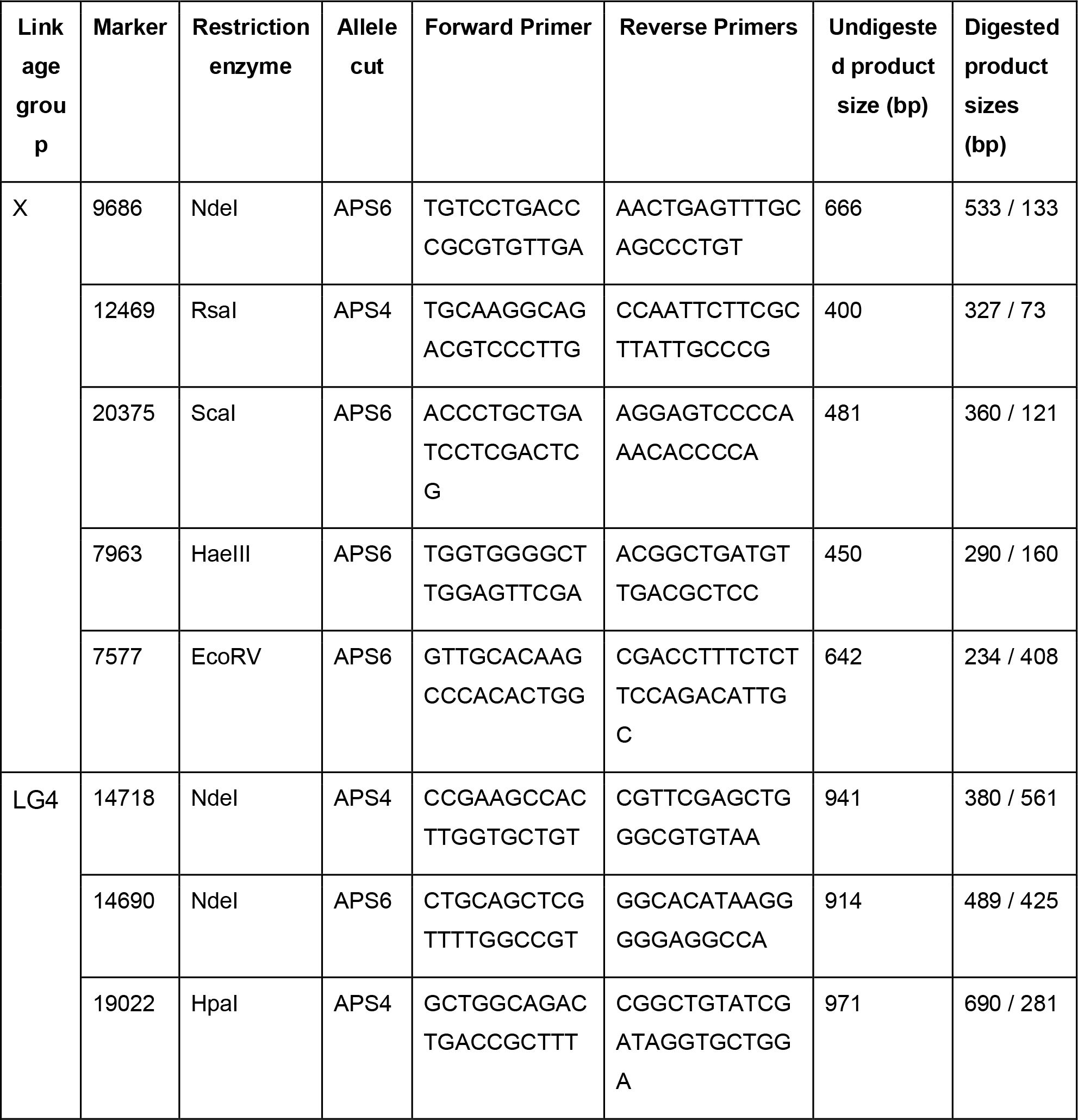
Summary of the markers used to genotype the X and LG4 chromosomes.

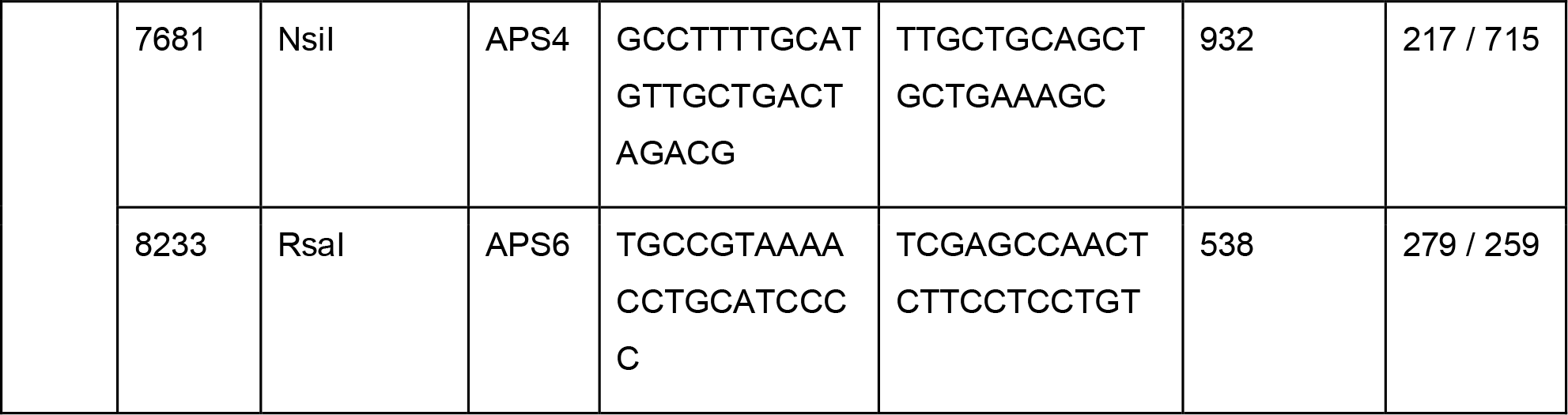

**Figure S1.**
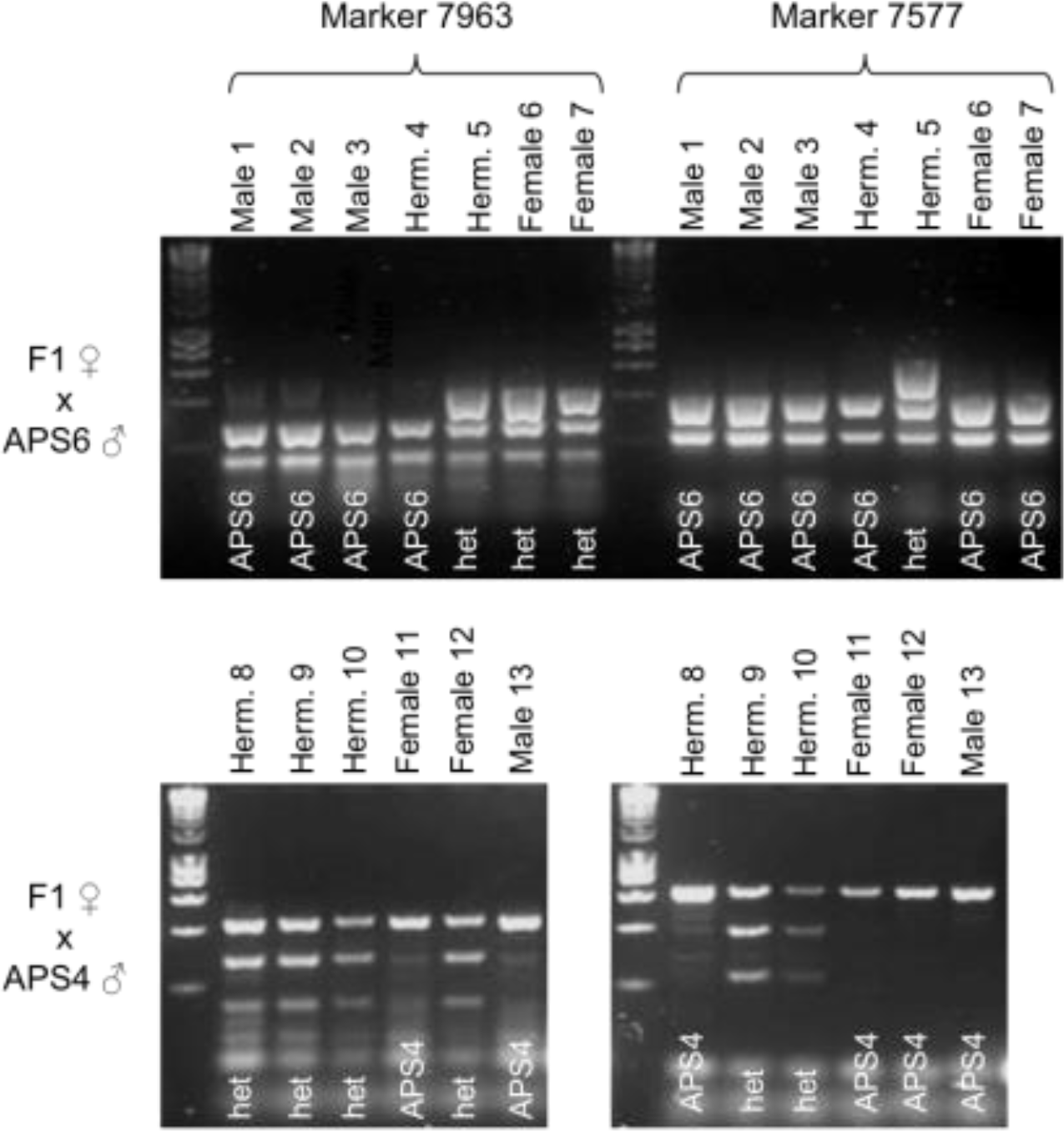
Example of X chromosome genotyping of F2s generated from crosses between F1 hybrid X_APS4_X_APS6_ females and either APS6 (upper panel) or APS4 (lower panel) males. From this genotyping, we can infer that some crossovers have occurred during female oogenesis as some F2 feminine do not display the same genotype across all the markers genotyped. The gel depicts only the two rightmost X markers (see figure 4A) as the crossovers were frequently observed between these markers (probably due to the subtelomeric position of the marker 7577). Genotypes are reported under the gel pictures. Numbers indicate individual animals.

**Figure S2.**
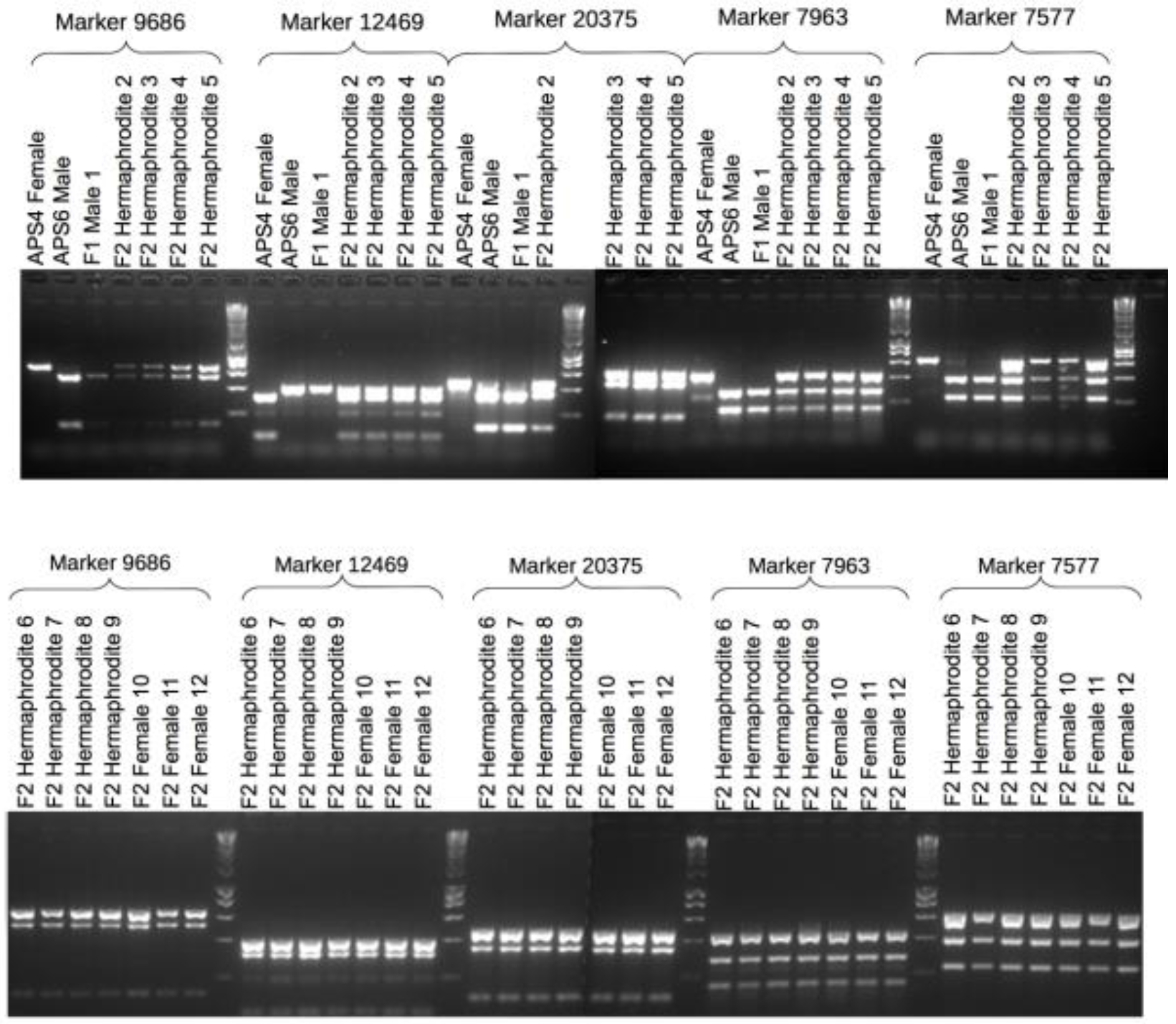
Example of X chromosome genotyping using amplification, digestion and electrophoresis of parental individuals and F2 females and hermaphrodites produced by selfing F1 hybrid hermaphrodites. F2 feminine progeny produced by hybrid F1 hermaphrodites are systematically heterozygous across the 5 X markers genotyped. Numbers indicate individual animals.

**Figure S3.**
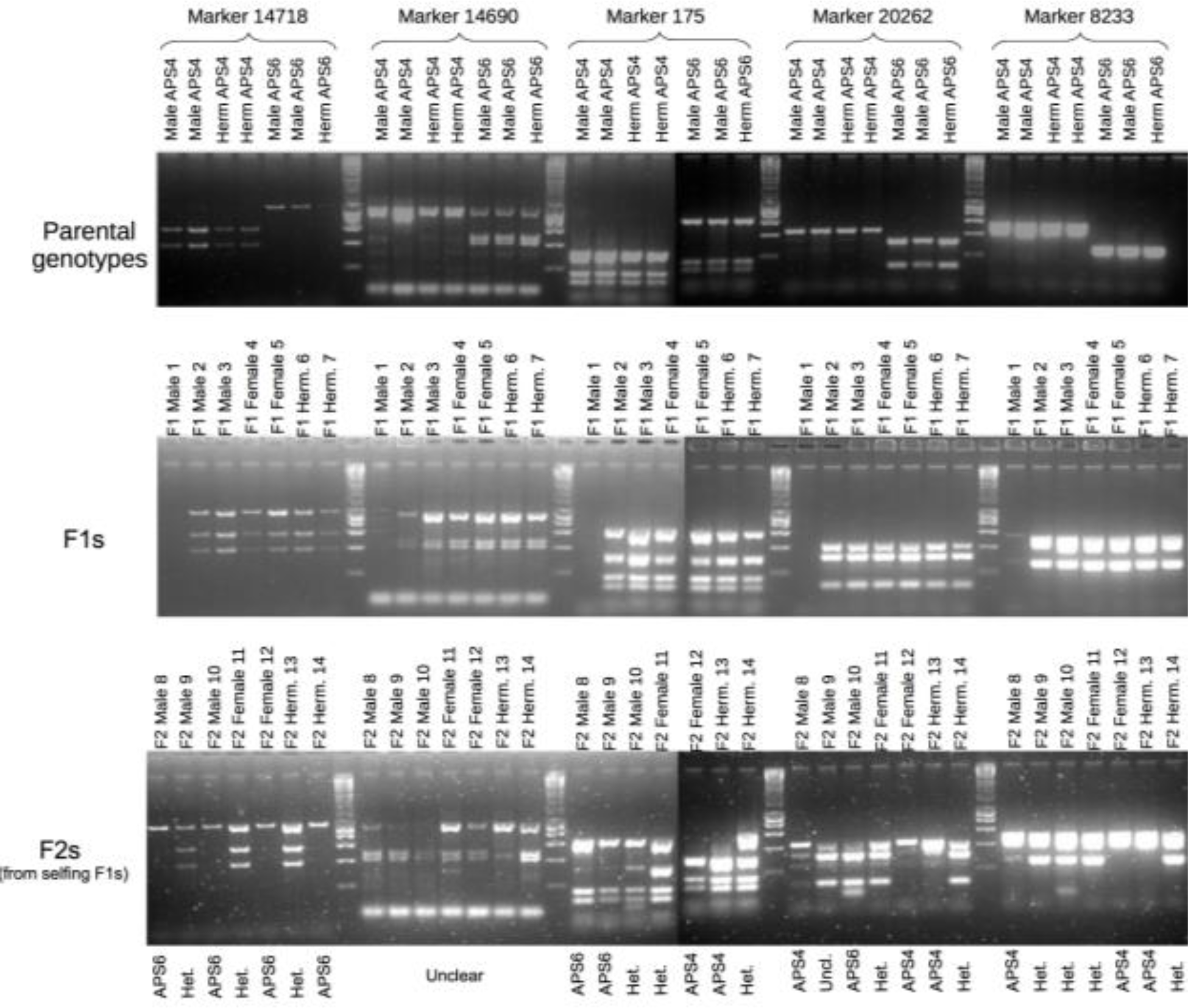
Example of LG4 (autosome) genotyping using amplification, digestion and electrophoresis. F2 genotypes are reported under the gel pictures. ‘Het.’ and ‘Uncl.’ stand for heterozygous and unclear genotypes, respectively. Numbers indicate individual animals.

## References

Albertson, D. G. & Thomson, J. N. 1993. Segregation of holocentric chromosomes at meiosis in the nematode, Caenorhabditis elegans. Chromosome Res, 1, 15–26.

Boyd, L., Guo, S., Levitan, D., Stinchcomb, D. T. & Kemphues, K. J. 1996. PAR-2 is asymmetrically distributed and promotes association of P granules and PAR-1 with the cortex in C. elegans embryos. Development, 122, 3075–84.

Brenner, S. 1974. The genetics of Caenorhabditis elegans. Genetics, 77, 71–94.

Chaudhuri, J., Bose, N., Tandonnet, S., Adams, S., Zuco, G., Kache, V., Parihar, M., Von Reuss, S. H., Schroeder, F. C. & Pires-Dasilva, A. 2015. Mating dynamics in a nematode with three sexes and its evolutionary implications. Sci Rep, 5, 17676.

Chaudhuri, J., Kache, V. & Pires-Dasilva, A. 2011. Regulation of sexual plasticity in a nematode that produces males, females, and hermaphrodites. Curr Biol, 21, 1548–51.

Church, D. L., Guan, K. L. & Lambie, E. J. 1995. Three genes of the MAP kinase cascade, mek-2, mpk-1/sur-1 and let-60 ras, are required for meiotic cell cycle progression in Caenorhabditis elegans. Development, 121, 2525–35.

Cortes, D. B., Mcnally, K. L., Mains, P. E. & Mcnally, F. J. 2015. The asymmetry of female meiosis reduces the frequency of inheritance of unpaired chromosomes. eLife, 4, 06056.

Dumont, J., Oegema, K. & Desai, A. 2010. A kinetochore-independent mechanism drives anaphase chromosome separation during acentrosomal meiosis. Nat Cell Biol, 12, 894–901.

Farrell, M. C. 2015. An investigation of hermaphroditism in R. sp. SB347. Master of Science, The College of William and Mary.

Félix, M. A. 2004. Alternative morphs and plasticity of vulval development in a rhabditid nematode species. Dev Genes Evol, 214, 55–63.

Golden, A., Sadler, P. L., Wallenfang, M. R., Schumacher, J. M., Hamill, D. R., Bates, G., Bowerman, B., Seydoux, G. & Shakes, D. C. 2000. Metaphase to anaphase (mat) transition-defective mutants in Caenorhabditis elegans. J Cell Biol, 151, 1469–82.

Hodgkin, J. 1979. Nondisjuncion mutants of the nematode Caenorhabditis elegans. Genetics, 91, 67–94.

Kanzaki, N., Kiontke, K., Tanaka, R., Hirooka, Y., Schwarz, A., Muller-Reichert, T., Chaudhuri, J. & Pires-Dasilva, A. 2017. Description of two three-gendered nematode species in the new genus Auanema (Rhabditina) that are models for reproductive mode evolution. Sci Rep, 7, 11135.

Kelly, W. G., Schaner, C. E., Dernburg, A. F., Lee, M. H., Kim, S. K., Villeneuve, A. M. & Reinke, V. 2002. X-chromosome silencing in the germline of C. elegans. Development, 129, 479–92.

Kiontke, K., Barriere, A., Kolotuev, I., Podbilewicz, B., Sommer, R., Fitch, D. H. & Félix, M. A. 2007. Trends, stasis, and drift in the evolution of nematode vulva development. Curr Biol, 17, 1925–37.

Macqueen, A. J., Phillips, C. M., Bhalla, N., Weiser, P., Villeneuve, A. M. & Dernburg, A. F. 2005. Chromosome sites play dual roles to establish homologous synapsis during meiosis in C. elegans. Cell, 123, 1037–50.

Mccaig, C. M., Lin, X., Farrell, M., Rehain-Bell, K. & Shakes, D. C. 2017. Germ cell cysts and simultaneous sperm and oocyte production in a hermaphroditic nematode. Dev Biol, 430, 362–373.

Mckee, B. D. & Handel, M. A. 1993. Sex chromosomes, recombination, and chromatin conformation. Chromosoma, 102, 71–80.

Meneely, P. M., Farago, A. F. & Kauffman, T. M. 2002. Crossover distribution and high interference for both the X chromosome and an autosome during oogenesis and spermatogenesis in Caenorhabditis elegans. Genetics, 162, 1169–77.

Nagaoka, S. I., Hassold, T. J. & Hunt, P. A. 2012. Human aneuploidy: mechanisms and new insights into an age-old problem. Nat Rev Genet, 13, 493–504.

Pasierbek, P., Jantsch, M., Melcher, M., Schleiffer, A., Schweizer, D. & Loidl, J. 2001. A Caenorhabditis elegans cohesion protein with functions in meiotic chromosome pairing and disjunction. Genes Dev, 15, 1349–60.

Phillips, C. M., Wong, C., Bhalla, N., Carlton, P. M., Weiser, P., Meneely, P. M. & Dernburg, A. F. 2005. HIM-8 binds to the X chromosome pairing center and mediates chromosome-specific meiotic synapsis. Cell, 123, 1051–63.

Rog, O. & Dernburg, A. F. 2013. Chromosome pairing and synapsis during Caenorhabditis elegans meiosis. Curr Opin Cell Biol, 25, 349–56.

Severson, A. F., Ling, L., Van Zuylen, V. & Meyer, B. J. 2009. The axial element protein HTP-3 promotes cohesin loading and meiotic axis assembly in C. elegans to implement the meiotic program of chromosome segregation. Genes Dev, 23, 1763–78.

Severson, A. F. & Meyer, B. J. 2014. Divergent kleisin subunits of cohesin specify mechanisms to tether and release meiotic chromosomes. eLife, 3, e03467.

Shakes, D. C., Neva, B. J., Huynh, H., Chaudhuri, J. & Pires-Dasilva, A. 2011. Asymmetric spermatocyte division as a mechanism for controlling sex ratios. Nat Commun, 2, 157.

Shakes, D. C., Wu, J. C., Sadler, P. L., Laprade, K., Moore, L. L., Noritake, A. & Chu, D. S. 2009. Spermatogenesis-specific features of the meiotic program in Caenorhabditis elegans. PLoS Genet, 5, e1000611.

Stiernagle, T. 2006. Maintenance of C. elegans. WormBook, 1–11.

Villeneuve, A. M. 1994. A cis-acting locus that promotes crossing over between X chromosomes in Caenorhabditis elegans. Genetics, 136, 887–902.

Winter, E. S., Schwarz, A., Fabig, G., Feldman, J. L., Pires-Dasilva, A., Muller-Reichert, T., Sadler, P. L. & Shakes, D. C. 2017. Cytoskeletal variations in an asymmetric cell division support diversity in nematode sperm size and sex ratios. Development, 144, 3253–3263.

Zickler, D. & Kleckner, N. 2015. Recombination, Pairing, and Synapsis of Homologs during Meiosis. Cold Spring Harb Perspect Biol, 7, a016626.

